# The Speed Limit of Visual Perception: Bidirectional influence of image memorability and processing speed on perceived duration and memory recall

**DOI:** 10.1101/2025.08.20.671225

**Authors:** Martin Wiener, Chloe Mondok, Alex Ma, Chetan Desai, April Joyner, Giuliana Macedo

**Author notes:** <u>Corresponding Author:</u> Martin Wiener, Department of Psychology, George Mason University, 4400 University Drive, Fairfax, VA 22030.

## Abstract

Visual stimuli are known to vary in their perceived duration, with some stimuli engendering so-called “time dilation” and others “time compression” effects. Previous theories have suggested these effects rely on the level of attention devoted to stimuli, magnitude of the stimulus dimension, or intensity of the population neural response, yet cannot account for the full range of experimental effects. Recently, we demonstrated that perceived time is affected by the image properties of scene clutter, size, and memorability (Ma, et al. 2024), with the former compressing and latter two dilating duration. Further, perceived duration also predicted recall of images 24h later, on top of memorability. To explain the memorability effect, we found that a recurrent convolutional neural network (rCNN) could recapitulate the time dilation effect by indexing the rate of entropy collapse, or “speed”, across successive timesteps, with more memorable stimuli associated with faster speeds. Here, we replicate and extend these findings via three experiments (n=20ea.) where subjects performed a sub-second temporal bisection task using memorability stimuli with increasing memorability, but a constant speed (exp 1), increasing speed, but constant memorability (exp 2), or increasing in both (exp 3), each followed by a surprise memory test 24hr later. We found that either increasing memorability or speed alone led to time dilation effects, with faster/slower speeds shifting memory recall by ∼8.4% in either direction. However, when both metrics increased, memorability dilated time while speed compressed it, while still improving recall overall. These findings can be explained by a model wherein the visual system is tuned to a preferred speed for processing stimuli that scales with the magnitude of visual response, such that stimuli closer to this speed are dilated in time. Overall, these findings provide a new lens for interpreting time dilation/compression effects and how visual stimuli are prioritized at temporal scales.

**Author Summary:** Recent work has demonstrated a link between perceived duration and memory recall, with memorable images dilating time, and perceived duration in turn affecting memory. The link between these two can be indexed by the speed at which images are processed in recurrent convolutional neural networks, yet the limits of this link remain untested. Here, we tested three groups on a timing task using three sets of stimuli selected according to both their memorability and speed. We find that when one factor is held constant, both memorability and speed still dilate perceived duration, with speed alone accounting for a shift in memory recall of ∼17%. However, when speeds are raised to their highest limits, perceived duration is instead compressed. An inverted-U shape rule between time dilation and compression over speed that is moderated by memorability explains this effect and suggests the visual system shifts between these modes to both gather more information and conserve energy, respectively.

## Introduction

The accurate and precise timing of sensory signals is an essential element of survival. Indeed, to be able to plan, anticipate, and properly react to stimuli in the environment, organisms are endowed with timing abilities that allow them to do so (Gibbon, et al. 1997). Notably, humans appear capable of adapting their sensory timing abilities to the temporal qualities of the environment they inhabit (Ossmy, et al. 2013; de Jong, et al. 2023) and further, other organisms appear to have evolved timing abilities relative to their ecological strata (Healy, et al. 2013). Yet, understanding how neural systems generate, adapt to, and accommodate timing signals remains underexplored.

Recently, Zheng and Meister (2025) have provided commentary on an issue related to the above: the rate at which information is processed. In their commentary, the authors surveyed a wide variety of human performance on various metrics, from problem solving to speech, and concluded that humans are bandwidth-limited in their behavior to 10 bits/second. The authors refer to this repeatedly as a “speed-limit”, with some speculation on its origin and purpose. While there has been some discussion generated from this commentary (see Sauerbrei & Pruszynski, 2025), one question remains to the use of the term “speed-limit” as a metaphor for the limits on human cognitive operations. That is, one can define a speed-limit under two frameworks: 1) it is an absolute limit on the maximum speed that can be achieved, such that higher speeds are impossible, or 2) it is an advisory limit on the fastest speed that can be maintained *safely*. Using the first definition, cognitive operations are unable to go faster than ∼10 bits/s simply because they cannot, whereas under the second definition, they *can* go faster but *shouldn’t* because there will be negative consequences to doing so. The obvious everyday analogy is one of driving speeds, where violating the speed limit can lead to accidents.

Related to the above, one can use the perception of time as a proxy for processing bandwidth. Notably, perceived duration is highly malleable, with numerous stimuli capable of engendering so-called “time dilation” and “time compression” effects, wherein stimuli can appear to last for subjectively longer and shorter amounts of time (Allman, et al. 2014; Matthews & Meck, 2018). Numerous stimuli have been found to alter perceived duration, such as stimulus size, brightness, frequency, contrast, color, motion, numerosity, and emotional content, among others. Recently, we (Ma, et al. 2024) found in a series of experiments that perceived duration was altered by features of scene images, including scene size, clutter, and memorability, with size and memorability dilating and clutter compressing perceived duration. Further, we found that the size of the time dilation effect with memorability was predictive of the ability of participants to recall the images 24 hours later.

Memorability itself is a metric calculated from memory recall performance and is suggested as an intrinsic feature of scene images (Isola, et al. 2013; Bainbridge, 2019). Further evidence suggests memorability is stable across individuals, is independent of attention, and is associated with a host of perceptual benefits (Khosla, et al. 2014; Goetschalckxn& Wagemans, 2019; Bainbridge, 2020). Yet, understanding what features drive memorability is still being disentangled, with a variety of high- and low-level features contributing (Isola, et al. 2013). Our previous report suggests that time – that is, the perceived duration – may be one feature contributing to memorability. In our previous report, to explain the effect of memorability, we processed the images used in our experiment (n=196) through a recurrent convolutional neural network (rCNN). The rCNN model is an extension of the well-known convolutional neural network (CNN) in which the model is allowed to iteratively process the presented image across a successive series of “timesteps” (Kietzman, et al. 2018; van Bergen & Kriegeskorte, 2020). The form of the iterative process can take different forms, such as feedback from higher layers to lower ones or local recurrence within a given layer (Nayebi, et al. 2018). We used an 8-layer rCNN network (BLnet; Spoerer, et al. 2020) in which each layer’s output was successively fed back to its input on the next time step, thus “unrolling” the network in time. This model processed each image across 8 successive timesteps, where each step represented a full forward-pass of the model. At the end of each forward-pass, a softmax layer provided a probability distribution of class labels, from which Shannon Entropy can be calculated as a proxy of model “certainty”; that is, the degree to which the model converged on a solution. Previous work with rCNNs has shown them to be superior at processing and classifying image categories, although the reasons for this superiority are still not well understood (Spoerer, et al. 2020). Further, the entropy of the model can be used as a proxy for the “speed” with which the model processes the image; previous work has shown that rCNN entropy is correlated with human reaction time (RT) data for image categorization and object recognition (Karapetian, et al. 2023; Sörensen, et al. 2023).

When we ran our images through BLnet and quantified entropy, we discovered that image memorability was associated with a faster “speed” with which the image converged on a solution. That is, entropy was found to decrease at a steeper rate for more memorable images. Further, by selecting an entropy threshold as a proxy for duration categorization (that is, whether a given arbitrary duration was considered “short” or “long), we observed that the model recapitulated the behavioral effect of memorability in humans and also predicted faster RTs for memorability judgments – which was observed. To describe the decline in entropy across successive timesteps, we fit the model with a simple, two-parameter power curve model

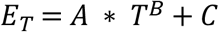

Where *E* represents the entropy at timestep *T*, and parameters *A, B*, and *C* reflect model free parameters. Crucially, the first parameter *A* determines the rate of decline in entropy and can be used as a proxy for model “speed”. As an additional finding, we note here that the faster speeds observed with higher memorability did not depend on the weights used for the model. That is, faster speeds for more memorable images were found with random weights. This finding, which accords with other findings that untrained CNN models with random weights are capable of detecting numerosity (Kim, et al. 2021), faces (Baek, et al. 2021) and music (Kim, et al. 2024), suggests that the decrease in entropy reflects intrinsic features of the images and not higher-order features that require image labels and model training.

Beyond the observed model findings, we applied the rCNN model to the entire LaMem dataset, consisting of 58,740 images (Khosla, et al. 2014). Here, we again observed that a significant relationship between memorability and model speed. Yet, importantly, we note here that the relationship exhibited marked heteroscedasticity; that is, as images increased in memorability, the spread of possible speeds grew wider. One implication of this is that the images selected in our previous study may have coincidentally been the right images for inducing time dilation; had we randomly sampled images that were at slower speeds for higher memorability, we would have concluded that model speed has no, or perhaps the opposite relationship with memorability. Further, we may then have found no behavioral effect or an opposite effect. This raises the general question of the relationship between memorability and model speed on perceived duration; that is, was the effect observed in our previous study entirely due to model speed, or memorability, or some combination of the two (and if so, which one has the larger influence?).

To answer the above questions, we conducted a new study where we sampled three new sets of images from the LaMem dataset according to memorability and the speed values calculated from the rCNN model and reported in our previous paper. In doing so, we sought to both replicate our previous findings and extend them to new images with different features, as well as further understand which features drive model speed and how they relate to both time perception and memory recall.

## Results

To begin, we recruited a sample of 60 participants, randomly assigning them to three separate groups. Each group was tested by a separate set of experimenters who were blind to the condition. Each group began by performing a temporal categorization (also known as bisection) task (see Methods). This task, which was identical to that used in our previous report (Ma, et al. 2024; Experiment 3), required participants to view image stimuli one-at-a-time for varying durations between 300 and 900ms, logarithmically-spaced and provide a speeded judgment regarding whether the interval presented was “short” or “long” based on all previously-presented intervals (Figure 1A). Previous research using this task has shown that subjects quickly adopt an internal standard within a few trials that remains reasonably stable throughout the session (Wearden & Ferrera, 1995; Wiener, et al. 2014).

**Figure 1.**
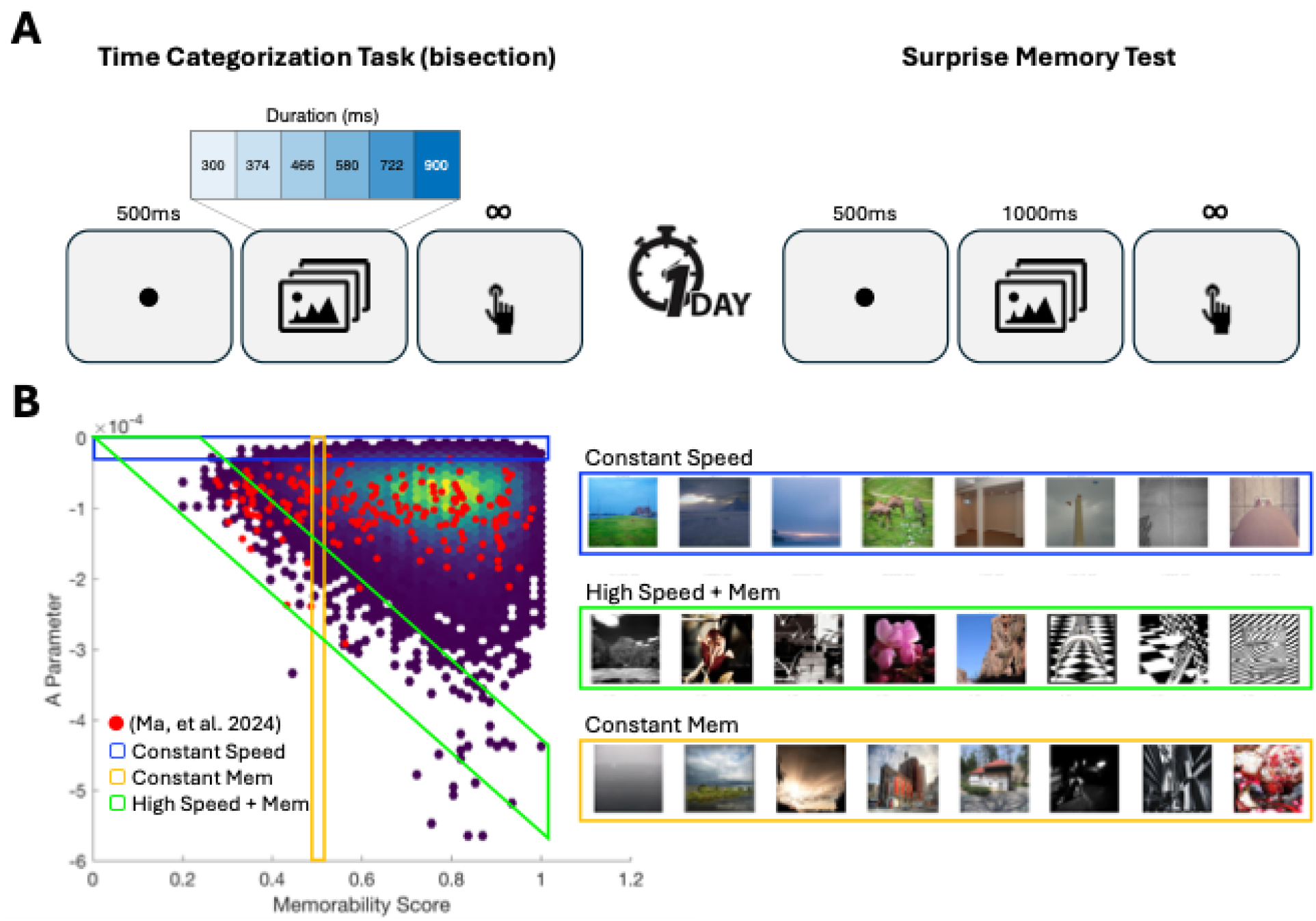
Task Design and Stimuli. **A**) Participants initially performed a Time Categorization (also called bisection) task in which a fixation point was followed by an image presented for one of 7 possible log-spaced durations between 300 and 900ms, which subjects then categorized as “short” or “long”. All participants returned the following day to perform a surprise memory test, in which stimuli from the previous day and a set of matched foils were presented with participants judging if they had seen the image before. **B**) Stimuli from the LaMem dataset with memorability and “speed” (as determined by the A parameter of our simple model). Red points indicate those stimuli used in our previous report (Ma, et al. 2024). Three new sets of stimuli were sampled from these images along three different axes: “Constant Speed” stimuli which increase in memorability yet maintain an approximately similar (slow) speed; “Constant Memorability” which increase in speed yet all have a memorability of 0.5; “High Speed + Memorability” which increase in memorability and all have respectively higher (fast) speeds. Examples of each set are shown on the right, which increase in either speed, memorability, or both from left-to-right− see Supporting Information for full stimuli.

Each group was presented with a different set of stimuli, consisting of 196 images (Figure 1B). These images were drawn from the LaMem dataset according to both the memorability score for that image and the value of the A parameter as determined by our simple model of rCNN entropy decrease (values are available at https://osf.io/fx3n2/). The first group, termed “Constant Speed” was selected by identifying those images across the range of memorability scores that had the correspondingly “slowest” speeds (i.e. the highest values of the A parameter). These images thus increased in memorability from 0.2 to 1 yet remained highly similar in terms of their speed. Qualitative observation of the stimuli showed them to typically consist of landscape images lacking in color, yet with little to distinguish the higher memorability images from the lower ones (Supporting Figure 1). The second group, termed “Constant Memorability” was selected by only sampling images with a memorability score of 0.5 (50% likely to be recalled), but with speeds that varied from the slowest (i.e. highest A value) to the fastest (i.e. lowest A value). These images varied from darker and cooler color tones and outdoor landscapes to urban and indoor scenes with a greater density of objects and artwork and variety of colors (Supporting Figure 2). The third group, termed “High Speed + Memorability” was selected in the opposite manner of the “Constant Speed” group by instead selecting those images across the range of memorability scores that had the correspondingly “fastest” speeds (i.e. lowest values of the A parameter). These images were highly stylized, with a mixture of color and black-and-white, high contrast, and with elements of graphic design (logos, symbols, artwork) (Supporting Figure 3). Subjects were given no instruction regarding the images themselves, only being told to attend to their duration.

Each participant was additionally asked to return one day later to complete a second session. In this session, participants were given a surprise memory test, where they were presented with the same 196 images as the previous day, along with 196 foil images sampled in the same manner (see Methods). Subjects were presented with each image and asked to indicate if they had seen it on the previous day.

For the first group (Constant Speed), a GLMM of response choices with memorability and speed as fixed effects and speed by subject as a random effect demonstrated a significant effect of memorability [χ^2^(1) = 69.097, *p*<0.001], speed [χ^2^(1) = 93.101, *p*<0.001], and an interaction between the two [χ^2^(1) = 211.628, *p*<0.001] (see Methods for details of the GLMM analysis). Fixed effects estimates revealed that both a higher memorability and a faster speed led to a higher likelihood of responding “long”,[Memorability: β = 1.242, SE = 0.149, *t* = 8.315, *p*<0.001; Speed: β = -0.284, SE = 0.029, *t* = -9.657, *p*<0.001] with a positive interaction between them [β = 1.291, SE = 0.089, *t* = 14.483, *p*<0.001]. Thus, the findings in this first group replicated both the previous effect of memorability and speed on perceived duration (Figure 2). On the second day, we also observed that images with higher memorability scores were associated with a greater accuracy for recall [*F*(6,114)=79.375, *p*<0.001, η^2^_p_ = 0.807] (Figure 3).

**Figure 2.**
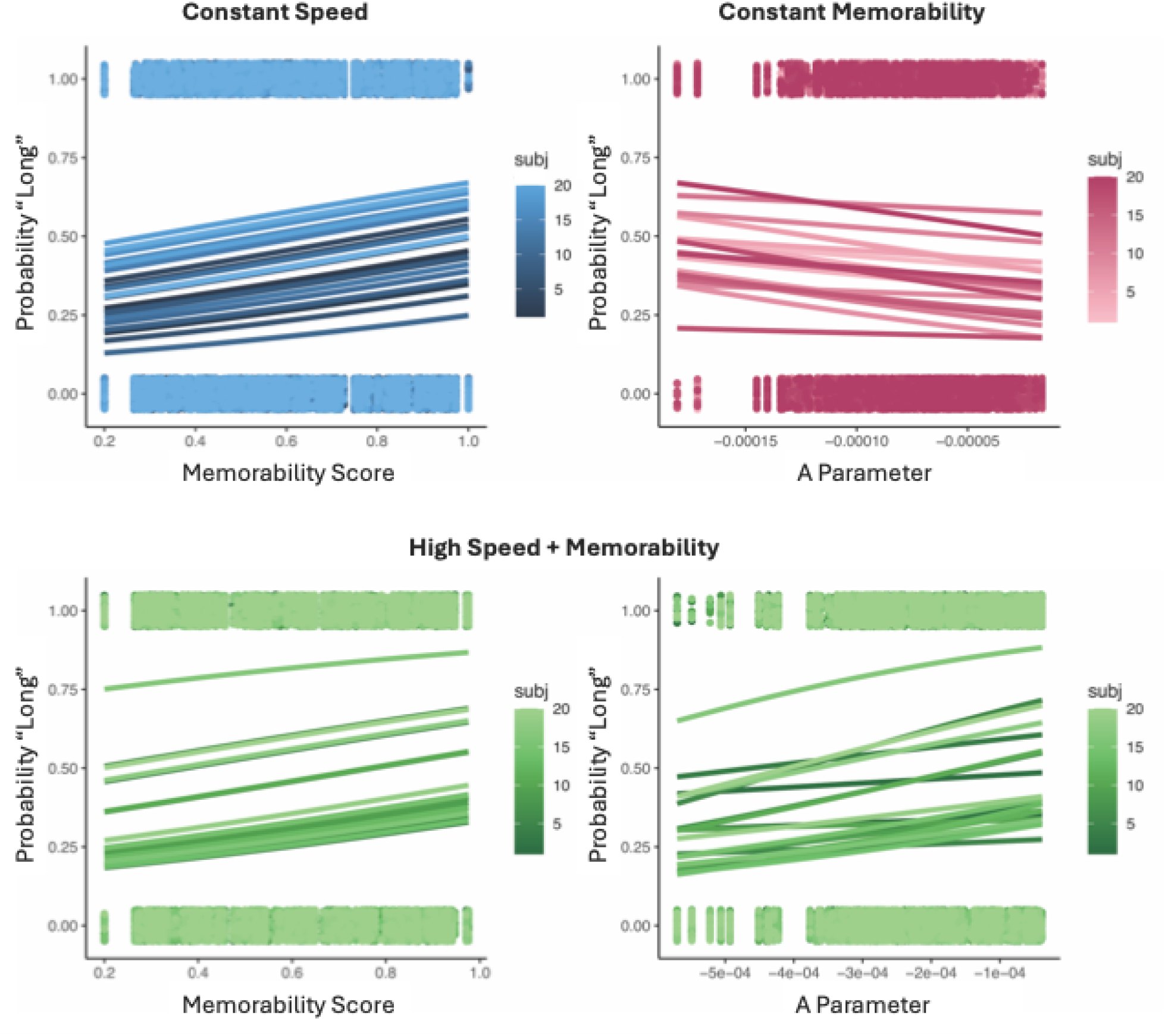
Separate and combined effects of speed and memorability on time estimates. For the either the Constant Speed or Constant Memorability groups, higher memorability stimuli or faster speeds (more negative A parameter values) led to an increase in the probability of responding “long”. In contrast, for the High Speed + Memorability group, higher memorability stimuli led to more “long” responses while faster speeds led the opposite effect. Plotted points represent individual trial responses for each subject, whereas smooth lines represent fits from our generalized linear model.

**Figure 3.**
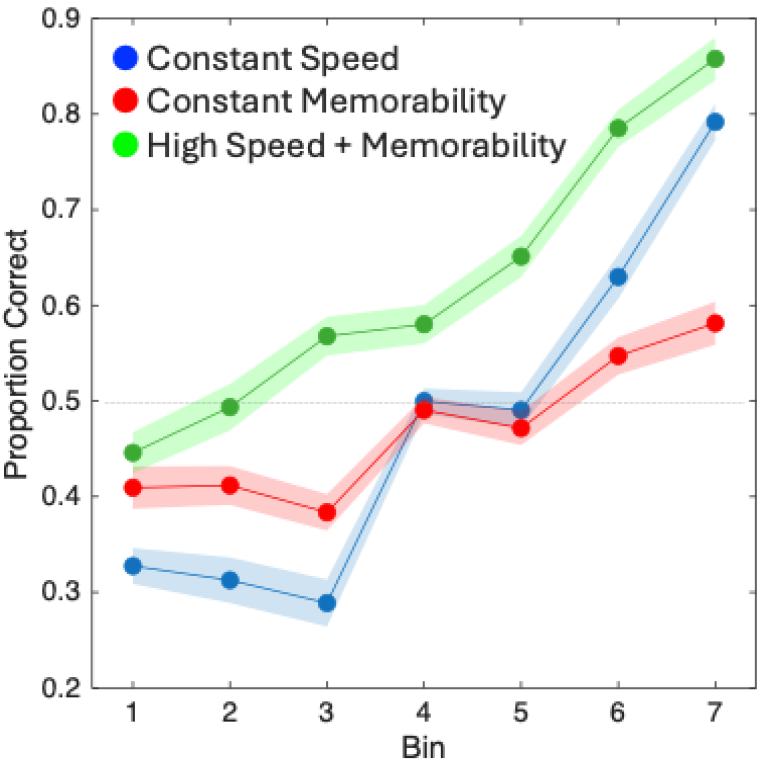
Memory recall performance on the surprise memory test. For the Constant Speed group, participants were more accurate for stimuli with higher memorability (higher bins). For the Constant Memorability group, where all stimuli had a memorability of 0.5, participants were more accurate at recalling stimuli with faster speeds (higher bins) and less accurate at recalling stimuli with slower speeds (lower bins). For the High Speed + Memorability group, participants were more accurate at recalling images that increased in both memorability and speed (higher bins). Further, accuracy in this group was significantly higher than either the Constant Speed or Constant Memorability groups. Plotted points represent average proportion correct with shaded regions representing within-subject standard error.

For the second group (Constant Memorability), a GLMM of response choices with speed as a fixed effect and subject as a random effect demonstrated a significant main effect of speed [χ^2^(1) = 58.025, *p*<0.001], such that higher speeds (i.e. more negative values of the A parameter) were associated with a higher probability of responding “long” [β = - 0.032, SE = 0.004, *t* = -7.623, *p*<0.001] (Figure 2). On the second day, we here observed that images with faster speeds were associated with a greater accuracy for recall [*F*(6,114)=13.097, *p*<0.001, η^2^_p_ = 0.408] (Figure 3). This finding is noteworthy, as the memorability for all these images was at 0.5, meaning participants should have been at chance in recalling them. Indeed, the mean hit rate across all images was 0.492, yet the spread of the mean hit rates across bins was 0.168, indicating that speed alone could increase or decrease the probability of recall on average by 8.4% in either direction.

For the third group (High Speed + Memorability), a GLMM of response choices with memorability and speed as fixed effects, as well as memorability and speed by subject as random effects demonstrated a significant main effect of memorability [χ^2^(1) = 47.607, *p*<0.001], speed [χ^2^(1) = 32.916, *p*<0.001], and an interaction between the two [χ^2^(1) = 10.279, *p*=0.001]. For the covariates, we observed positive effect of memorability, such that higher memorability images were again associated with more “long” responses [β = 2.283, SE = 0.217, *t* = 10.506, *p*<0.001]; however, for speed we here observed a positive effect [β = 0.034, SE = 0.006, *t* = 6.098, *p*<0.001], indicating that *slower* speeds were associated with a higher probability of responding “long”, in contrast to the results from the Constant Memorability and Constant Speed groups (Figure 2). The interaction effect was also positive [β = 0.023, SE = 0.007, *t* = 3.066, *p*=0.002]. Despite this difference, on the second day, we again found that higher memorability images with faster speeds were associated with greater accuracy for recall [*F*(6,114)=44.005, *p*<0.001, η^2^_p_ = 0.698] (Figure 3).

To determine if there were any differences between the image sets on the recall day, an omnibus ANOVA with all three groups found a significant main effect of group [*F*(2,57)=5.836, *p*=0.005, η^2^_p_ = 0.17] and interaction between memorability/speed bin and group [*F*(12,342)=10.343, *p*<0.001, η^2^_p_ = 0.266]. Post-hoc tests with Bonferroni correction found that the High Speed + Memorability group exhibited higher recall than both the Constant Memorability [*t* = 2.743, *p* (corrected) = 0.024, Cohen’s *D* = 0.779] and the Constant Speed [*t* = 3.136, *p* (corrected) = 0.008, Cohen’s *D* = 0.891] groups. Notably, the mean difference between the High Speed + Memorability group and the other two was 0.178 and 0.155, respectively, which is similar to the shift induced by speed in the Constant Memorability group (0.168), thus suggesting a general effect of speed on memory recall in this range.

The difference in effect direction between the High Speed + Memorability group and the Constant Memorability and Constant Speeds group initially presented a puzzle, as we had expected that higher speeds would induce a larger time dilation effect than in either group, whereas here observed a time compression effect. To determine the reason for this difference, we examined responses within this group more closely as a function of both memorability and speed. To do this, we divided memorability and speed for this group into seven separate bins and calculated the mean proportion of “long” responses as a function of each. Here, we observed that, within each memorability bin, higher speeds were indeed associated with a decrease in responding “long”; however, the rate of this decrease varied as a function of the memorability bin itself, such that higher memorability was associated with a shallower time compression effect across speed bins (Figure 4). For a broader view, we combined the data from the High Speed + Memorability and Constant Memorability groups; by doing so, we obtained a full set of speeds that covered the entire set of images where memorability was approximately 0.5. Here, we divided speed into 11 bins due to the wider coverage. We observed that the proportion of “long” responses initially increased with higher speeds – a time dilation effect – before inflecting and shifting downward into a time compression effect (Figure 4). This effect could easily be described as a quadratic effect with a concave shape.

**Figure 4.**
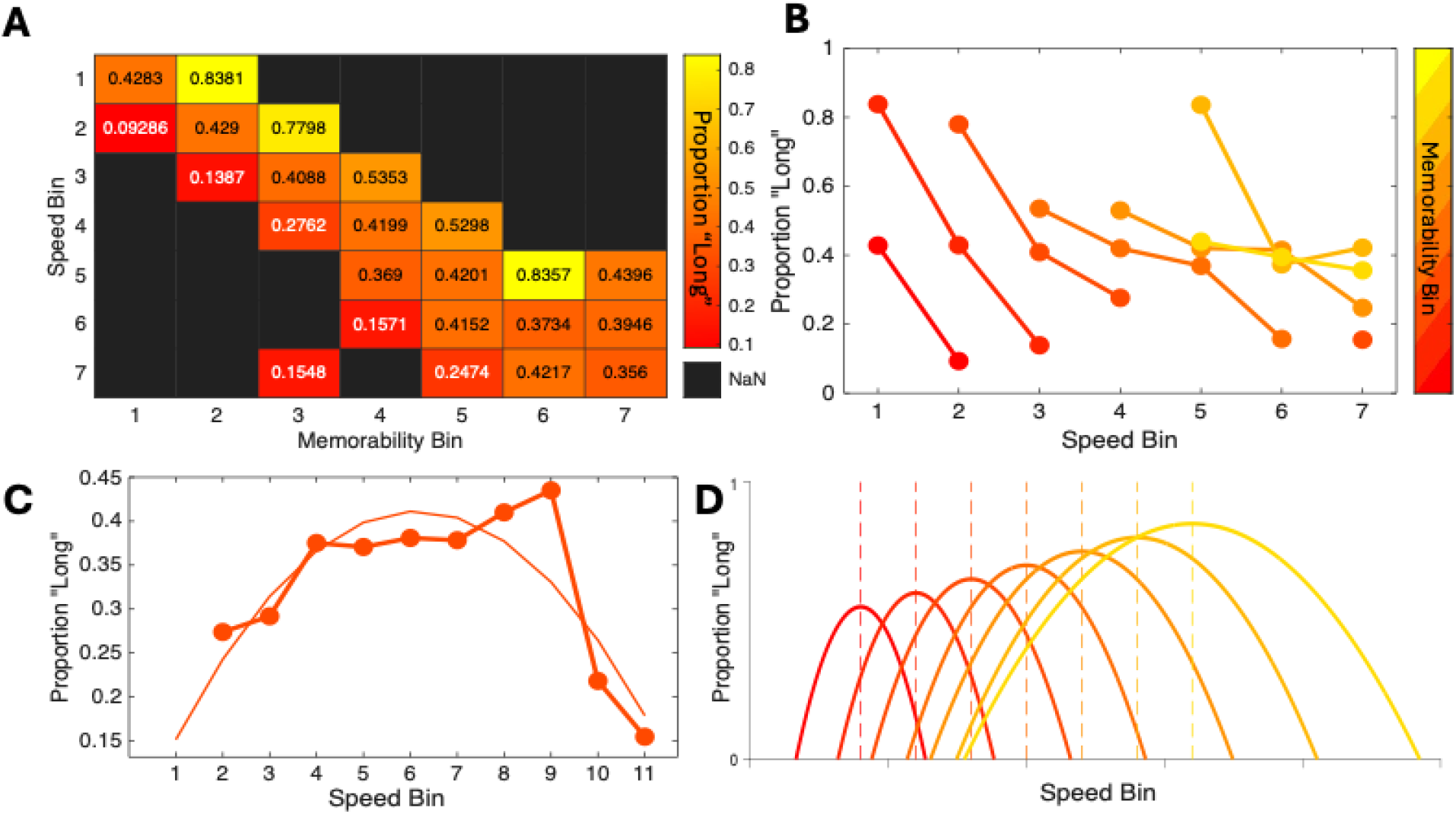
Relationship between speed and memorability. **A**) Heatmap of mean proportion of “long” responses for the High Speed + Memorability group as a function of both memorability and speed, each of which were separated into seven equally-spaced bins. **B**) Same data as in (A) but presented as a line plot with each line representing a successive memorability bin. Within each memorability bin, faster speeds are associated with lower proportions of responding “long”, yet this effect becomes shallower as memorability increases; note also that higher memorability bins are associated with higher average proportions of “long” responses in general. **C**) Mean proportion “long” but for data combined from the High Speed + Memorability and Constant Memorability groups; these data comprise images at ∼0.5 memorability, but across 11 speed bins. Here, the results follow an inverted-U shape with faster speeds leading to progressively higher proportions of responding “long” followed by an inflection− overlaid curve represents a quadratic fit.

Taking the above observations into account, a simple rule can be derived to explain the opposing effects between low and high speeds and time dilation and compression. First, dilation and compression follow an inverted-U, quadratic-like trend with increasing speed. The precise shape of this curve exhibits a leftward-skew, which could either be described by a skewed Gaussian or by an exponential quadratic function. In either case, three aspects of this curve are affected by memorability: 1) higher values of memorability lead to wider spreads of the curve; by increasing the width, we account for the more gradual decrease in the proportion of “long” responses observed. 2) Higher values of memorability displace the peak of this function to higher speeds; this can account for how the same speed value for a given image can induce a higher/lower time dilation effect. 3) Higher values of memorability increase the peak height of the function; this accounts for the general time dilation effect that memorability has, while still accommodating the more gradual time compression effect at high speeds. Altogether, we can account for the shift from time dilation to compression using a single scalar: memorability. The resulting quadratic function is thus

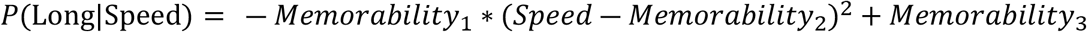

The initial value of Memorability(1) leads to progressively wider spreads, while the second value of Memorability(2) displaces the vertex of the curve to the right and the third value of Memorability(3) increases the vertex height (Figure 4). If needed, covariates can be added to each memorability value to change the shape as appropriate. We stress here that this model is meant to be merely illustrative of the proposed, inverted-U shaped form the relationship between speed and perceived duration takes. We turn to further discussion of this relationship below.

## Discussion

The results of above series of experiments both replicate and extend our previous findings. That is, we demonstrate here again that scene images with higher memorability scores are perceived for longer amounts of time than those with lower memorability scores. However, the findings as they relate to “speed” – as determined by the rate of entropy decline in a rCNN model – provide a more complicated relationship with perceived duration. While the results may appear contradictory at first, they in fact provide more holistic view of how the brain integrates visual information into time estimates.

First, our results demonstrate that when rCNN model speed is kept at an approximately constant and slow rate, the memorability of scene images is still able to induce time dilation effects. This finding is notable as the stimuli employed tended to be more colorless, flat, low in contrast and minimalist, and so dilation effects likely resulted from other features of the images; indeed, the more memorable images tended to be more unusual, artistic, or have human-like features. Yet, many of the more memorable images also had little to distinguish them from the less memorable ones, suggesting there were still as-yet-undetermined features within these images that led to time dilation effects. With regards to speed, as these images were all from the slower range of possible speeds, the lack of color and detail suggest that rCNN model speed is in-part determined by these features. Indeed, we found that selected images with much higher speeds tended to be more colorful, contrastive, and edgier. We additionally verified the increase in memorability for these images on the surprise memory test one day later, where more memorable images were more likely to be recalled. Despite this, however, we do note that the overall memorability of these images was lower than expected. Our sampling strategy was to select images with memorability from 0.2 to 1 in seven bins, yet generally the actual recall performance fell below the average memorability of each bin.

Second, we observed that when memorability was held constant at 0.5, yet speed was allowed to vary, faster speeds were associated with a time dilation effect. Here the images progressed from outdoor landscapes and nighttime images to indoor and urban environments with more colors, to artwork and artificial designs. This finding thus supports the idea that memorability and rCNN speed are distinct qualities of images, with each having an influence on perceived duration. Further, we also observed that model speed affected memory recall performance the following day. Here, as all images were drawn from the 0.5 memorability pool, the prediction is that participants should be at chance in recalling them. Instead, we found that subjects were worse at recalling images with slower speeds and better at recalling images with faster speeds, with a mean difference of ∼17%. This finding suggests that speed, and in-turn differences in perceived duration, can account for shifts in memory performance to this extent. We note that a shift of this size is similar to our previous report, where reproduced duration accounted for memory recall performance beyond the memorability score for an image (Ma, et al. 2024).

Finally, we observed that when images were selected with very fast speeds as memorability increased, these images led to a time dilation effect with memorability yet a time compression effect with speed. While the result appeared contrary to the previous two image sets, it is important to stress that the speeds employed here were very different from the previous two sets, in that they were all among the fastest speeds for each memorability bin. Thus, while the speeds covered some of the same area as the Constant Memorability group, they were associated with images of very different memorability. Upon closer inspection, we observed that the time compression effect was locally affected by the memorability bin; that is, within each memorability bin we observed a time compression effect, but this effect was modulated by the memorability itself. Specifically, we found that higher memorability was associated with a shallower time compression effect. When we combined these data with those of the Constant Memorability group, thus encompassing the full range of speeds in the ∼0.5 memorability bin, we observed an inverted-U shape effect on perceived duration, with faster speeds initially associated with time dilation before inflecting into a time compression effect at the highest speeds. To account for the inverted-U shape, as well as the general increase in time dilation with memorability and the shallower time compression effect with memorability, we calculated a simple quadratic rule where the effect of speed on perceived duration depends on the memorability of the stimulus; higher memorability shifts the speed at which time dilation inflects to time compression to a faster rate.

Altogether, the above finding of an inverted-U effect of speed suggests that this measure is bound by memorability. In terms of neural plausibility, we note that memorability has been linked to both neuronal population magnitude, where neurons in inferotemporal cortex fire more vigorously to stimuli higher in memorability, as well as population variance, where these neurons fire more consistently to more memorable stimuli (Jaegle, et al. 2019). While it is unknown what the neural correlates of faster speeds are, we speculate here that model speed is driven in part by the fidelity with which the image aligns with the convolutional filters at each layer of the network. As each image is iterated, any image that contains intrinsic features that align with the filters (e.g. clearer edges) will lead to a stronger response in the softmax layer, and thus a decrease in entropy. We further suggest here that, at the neural level, a larger, more consistent neural response can accommodate a faster speed stimulus.

Yet, why might this rule exist at all? Even with higher memorability accommodating greater speeds, why should there be an inflection from time dilation to time compression? We here suggest that the visual system (and indeed, perhaps all sensory systems) are tuned to a specific duration range when encountering complex stimuli. That is, to maintain the present perceptual moment, one cannot experience time as too fast or too slow, as the timing between events would therefore become too unpredictable for accurately timed behavior. Therefore, time dilation and compression effects may exist to maintain temporal stability. Indeed, other visual domains have demonstrated balances between time dilation and compression, although not with this interpretation. For example, recent work has shown that eye-blinks while timing visual stimuli lead to time compression (Grossman, et al. 2019), whereas stimuli that are presented several seconds after an eye-blink are dilated (Terhune, et al. 2016). Similarly, in the well-known chronostasis phenomenon, where saccades lead to a time dilation effect, recent work has shown a comparative time compression effect afterwards (Chen, et al. 2025). Also, in the similarly well-described number-time illusion, where larger numerals are perceived for longer intervals than shorter intervals, a contrastive after-effect has been shown such that larger numerals on the previous trial lead to shorter perceived intervals on the present one (Wehrman, et al. 2020). In our own previous report on memorability, we also observed that scene images depicting larger sizes were dilated, whereas those that were more cluttered were compressed. In the context of our present results, these effects can be explained by suggesting that the brain uses time to gather more information about a stimulus based on its features, yet when those features are too complex or overwhelming it compresses time to conserve energy and maintain perceptual stability. The point at which dilation turns to compression is thus a function of the energetic cost of processing the stimulus, which here is indexed by image memorability. To follow the “speed limit” analogy used in the introduction and by Zheng & Meister (2025), this would be more similar to an advisory limit; the visual system *can* perceive things at higher rates, but doing so entails a cost that is curtailed (i.e. by slowing down when traveling too fast). Further, just as the speed limit is faster on roads that accommodate higher speeds (i.e. a highway versus a local road), the visual system can accommodate dilation for stimuli at higher speeds if the neural population response is sufficiently large. These findings provide a new lens for time dilation and compression effects, putting both in the domain of an information-gathering strategy.

One final point remains to be explained – that regarding the memory recall performance on the second day. Despite the time compression effect observed with higher speeds, these stimuli were overall better recalled than those with slower speeds. Further, for the stimuli in the Constant Memorability group, these had all been sampled from those with memorability scores of 0.5, meaning previous participants had found them at chance for recall, yet our participants varied by a significant amount; if previous participants recalled them at chance, why did our group vary, and why weren’t the numbers in the LaMem dataset the same as ours? We suggest that it is unlikely the LaMem numbers were wrong, given the scores come from a large number of subjects, and further the split-half reliability is high (Khosla, et al. 2014). Instead, we suggest the differences between our participant memory recall scores and LaMem are due to context effects. Specifically, the images presented to subjects in LaMem varied widely in memorability and speed, whereas the stimuli presented to subjects here were in a narrow range. Indeed, a faster speed image with a predicted memorability of 0.5 may be more memorable when it is presented in the context of other 0.5 memorability images with slower speeds. Likewise, a set of images that all have slower speeds may be less memorable overall, whereas a set of images that all have high speeds may be more memorable overall. Previous work has similarly shown that image memorability is affected by local and global context; that is, whether the images are all memorable or forgettable (Gedvilla, et al. 2023) or whether the previous trial image was memorable or forgettable (Bainbridge, 2025). For example, among many memorable images, a forgettable one may become more memorable by the context, or conversely a memorable image may become forgettable if surrounded by forgettable images. Testing this possibility, as well as if speed is similarly affected, remains for future work.

### Time and Speed as Information Gathering

The results of the above experiments both replicate and extend our previous finding. Further, they add an important constraint on models of how the encoding of visual information affects perceived duration. Under this constraint, the brain is capable of processing images associated with faster rCNN model speed by dilating time, yet shifts to compression when those images are overly complicated or stimulating. Future work will need to determine how faster speeds alter neural processing, whether by affecting population magnitude or variance. A further prediction of this work is that specific stimuli could be generated to alter perceived duration, by taking memorability and speed into account, to induce time dilation or compression. Likewise, just as algorithms exist that can alter the memorability of a given image, via generative adversarial networks (Goetshalckx, et al. 2019), one could use a similar process to alter the speed of a given image, and so alter perceived duration and – indirectly – memorability. That is, one could take a memorable image, and make it even more, or less memorable. Future work will be necessary to test these possibilities.

## Materials and Methods

### Participants

A total of 60 participants [Mean Age 19.61 ± 2.49; 40 females] performed the experiment, randomly assigned to three separate groups of 20; further, each group of subjects was tested by a different set of experimenters, who were blind to the group they were testing. All participants were undergraduates at George Mason University and were compensated via course credit. All subjects were additionally right-handed, psychologically and neurologically healthy, and had normal or corrected-to-normal vision.

### Ethics Statement

Written consent was provided by each participant, and the protocol was approved by the University Institutional Review Board.

### Procedure

All participants performed a temporal categorization (also known as bisection) task with sub-second stimuli. In this task, each trial began with a central fixation point for 500ms, followed by a central image presented for one of seven durations logarithmically-spaced between 300 and 900ms (Figure 1). Following the image, subjects were instructed to respond as quickly yet as accurately as possible whether the duration of the image presented was “long” or “short”, by pressing the ‘S’ and ‘L’ keys, respectively. Participants were told to initially guess for the first few trials and then make their judgments according to the average of all durations they have seen. No specific instructions were given regarding the images. Images were presented in a series of 196 trials per block, with a break in between each block, for a total of 7 blocks and 1372 trials. The task design is identical to that used in our previous report (Ma, et al. 2024; Experiment 3).

After performing the categorization task, all participants were asked to return the next day for a follow-up session. No details regarding the second session was given. For this session, participants were presented with a surprise memory test. In this task, each trial began with a central fixation point for 500ms, followed by a central image (0.5 x 0.5 in PsychoPy height units) presented for 1000ms. Participants were instructed to indicate if the image presented was one they had seen on the previous day, by pressing the ‘Y’ key if they had, and the ‘N’ key if they hadn’t. A total of 392 trials were presented in a single block, consisting of 196 images and 196 foils (see below). The task design is identical to that used in our previous report (Ma, et al. 2024; Experiment 4).

All experiments were conducted in a testing room with participants seated at a comfortable distance from a 27” Gaming Monitor running at 100Hz. All responses were collected on a mechanical keyboard with a polling rate of 1000Hz. The experiments were programmed using PsychoPy (Peirce, 2007).

### Stimuli

Each group of participants performed the tasks as described above but using distinct sets of stimuli. The first set, referred to as “Constant Speed”, was selected from the LaMem dataset (http://memorability.csail.mit.edu/) by identifying images that increased in memorability yet maintained a constant “speed”, as characterized by the A parameter of the simple entropy collapse model in our previous report (Ma, et al. 2024). To accomplish this, images were sampled by separating all memorability scores into 7 equally spaced bins and, within each bin, sorting images by the A parameter and selecting the images with the highest A parameter. This process was repeated equally across each bin until a total of 392 images were selected; these images were then randomly split into two sets of 196 images each to be used as test and foil images as described above (Average A parameter = -2.36e-05 ± -1.72e-05). We note that the original set from which all values were drawn are available at (https://osf.io/fx3n2/). Full lists of stimuli used are also available at (https://osf.io/5bfvh/).

For the second set, referred to as “Constant Memorability”, the same process as above was conducted except by only selecting memorability images with a memorability score of 0.5. The A parameter values additionally separated into 7 equally spaced bins in a similar manner to that described above, again resulting in 392 images that were split into two sets.

For the third set, referred to as “High Speed + Memorability”, the process was the same as for the “Constant Speed” set, except that the lowest (most negative) A parameter values were chosen for each memorability bin, again resulting in 392 images that were randomly split into two sets.

### Analysis

Each participants data were initially filtered by removing outlier trials, defined as those with a reaction time greater than 3 standard deviations from the mean of the log-transformed distribution of reaction times. For each group, we employed a generalized linear mixed model (GLMM) analysis. All analyses were conducted in R using the lme4 and easystats package (Bates, et al. 2015; Lüdecke, et al. 2022). A binomial distribution was used with a logit link function. For random effects, we opted for a balance between the “keep it maximal” strategy (Barr, et al. 2013) and a parsimonious one (Bates, et al. 2018); that is, we initially fit each model with all the slopes of each covariate set as random effects by subject and piecewise removed them until a model was retained where 1) the model converged, 2) the model fit was not singular, 3) the residuals were normally distributed and none of the random effect variances were at zero (Scandola & Tidoni, et al. 2024). Model comparisons were conducted via nested Chi-Squared Likelihood tests using Type III Sum of Squares.

For the analysis of the second session, the hit rate (proportion correct for images presented on the previous day) was calculated for images within each of the seven bins used in our sampling strategy. A repeated-measures ANOVA was then conducted on these values separately for each group. An omnibus mixed-model ANOVA was also conducted comparing hit rate across bins and between groups.

## Acknowledgements

We thank numerous attendees at the Vision Sciences Society (VSS) 2025 annual meeting for helpful comments on these findings, which were presented in a talk session. The authors also wish to thank Bryce Sullivan for his assistance with this project.

## Supporting Information

Behavioral data for this project are available at: https://osf.io/5bfvh/

## Notes

### Competing Interest Statement

The authors have declared no competing interest.

## References

Allman, M. J., Teki, S., Griffiths, T. D., & Meck, W. H. (2014). Properties of the internal clock: first-and second-order principles of subjective time. Annual review of psychology, 65, 743–771.

Bainbridge, W. A. (2019). Memorability: How what we see influences what we remember. In Psychology of learning and motivation (Vol. 70, pp. 1–27). Academic Press.

Bainbridge, W. A. (2020). The resiliency of image memorability: A predictor of memory separate from attention and priming. Neuropsychologia, 141, 107408.

Bainbridge, W. (2025). What divides and unites our memories: multi-factor trial-wise predictions of memory across 6+ million trials. PsyArXiv 10.31234/osf.io/a79h6_v1

Chen, L., Grzeczkowski, L., Müller, H. J., & Shi, Z. (2025). Saccade-induced temporal distortion: opposing effects of time expansion and compression. Psychological Research, 89(2), 86.

de Jong, J., van Rijn, H., & Akyürek, E. G. (2023). Adaptive encoding speed in working memory. Psychological Science, 34(7), 822–833.

Gedvila, M., Ongchoco, J. D. K., & Bainbridge, W. A. (2023). Memorable beginnings, but forgettable endings: Intrinsic memorability alters our subjective experience of time. Visual cognition, 31(5), 380–389.

Gibbon, J., Malapani, C., Dale, C. L., & Gallistel, C. R. (1997). Toward a neurobiology of temporal cognition: advances and challenges. Current opinion in neurobiology, 7(2), 170–184.

Goetschalckx, L., & Wagemans, J. (2019). MemCat: a new category-based image set quantified on memorability. PeerJ, 7, e8169.

Goetschalckx, L., Andonian, A., Oliva, A., & Isola, P. (2019). Ganalyze: Toward visual definitions of cognitive image properties. In Proceedings of the ieee/cvf international conference on computer vision (pp. 5744-5753).

Grossman, S., Gueta, C., Pesin, S., Malach, R., & Landau, A. N. (2019). Where does time go when you blink?. Psychological science, 30(6), 907–916.

Healy, K., McNally, L., Ruxton, G. D., Cooper, N., & Jackson, A. L. (2013). Metabolic rate and body size are linked with perception of temporal information. Animal behaviour, 86(4), 685–696.

Isola, P., Xiao, J., Parikh, D., Torralba, A., & Oliva, A. (2013). What makes a photograph memorable?. IEEE transactions on pattern analysis and machine intelligence, 36(7), 1469–1482.

Jaegle, A., Mehrpour, V., Mohsenzadeh, Y., Meyer, T., Oliva, A., & Rust, N. (2019). Population response magnitude variation in inferotemporal cortex predicts image memorability. Elife, 8, e47596.

Karapetian, A., Boyanova, A., Pandaram, M., Obermayer, K., Kietzmann, T. C., & Cichy, R. M. (2023). Empirically identifying and computationally modeling the brain–behavior relationship for human scene categorization. Journal of cognitive neuroscience, 35(11), 1879–1897.

Khosla, A., Raju, A. S., Torralba, A., & Oliva, A. (2015). Understanding and predicting image memorability at a large scale. In Proceedings of the IEEE international conference on computer vision (pp. 2390-2398).

Kietzmann, T. C., Spoerer, C. J., Sörensen, L. K., Cichy, R. M., Hauk, O., & Kriegeskorte, N. (2019). Recurrence is required to capture the representational dynamics of the human visual system. Proceedings of the National Academy of Sciences, 116(43), 21854–21863.

Ma, A. C., Cameron, A. D., & Wiener, M. (2024). Memorability shapes perceived time (and vice versa). Nature Human Behaviour, 8(7), 1296–1308.

Matthews, W. J., & Meck, W. H. (2016). Temporal cognition: Connecting subjective time to perception, attention, and memory. Psychological bulletin, 142(8), 865.

Nayebi, A., Bear, D., Kubilius, J., Kar, K., Ganguli, S., Sussillo, D., … & Yamins, D. L. (2018). Task-driven convolutional recurrent models of the visual system. Advances in neural information processing systems, 31.

Ossmy, O., Moran, R., Pfeffer, T., Tsetsos, K., Usher, M., & Donner, T. H. (2013). The timescale of perceptual evidence integration can be adapted to the environment. Current biology, 23(11), 981–986.

Sauerbrei, B. A., & Pruszynski, J. A. (2025). The brain works at more than 10 bits per second. Nature Neuroscience, 1-2.

Sörensen, L. K., Bohté, S. M., De Jong, D., Slagter, H. A., & Scholte, H. S. (2023). Mechanisms of human dynamic object recognition revealed by sequential deep neural networks. PLOS Computational Biology, 19(6), e1011169.

Spoerer, C. J., Kietzmann, T. C., Mehrer, J., Charest, I., & Kriegeskorte, N. (2020). Recurrent neural networks can explain flexible trading of speed and accuracy in biological vision. PLoS computational biology, 16(10), e1008215.

Terhune, D. B., Sullivan, J. G., & Simola, J. M. (2016). Time dilates after spontaneous blinking. Current Biology, 26(11), R459–R460.

van Bergen, R. S., & Kriegeskorte, N. (2020). Going in circles is the way forward: the role of recurrence in visual inference. Current Opinion in Neurobiology, 65, 176–193.

Wehrman, J. J., Kaplan, D. M., & Sowman, P. F. (2020). Local context effects in the magnitude-duration illusion: Size but not numerical value sequentially alters perceived duration. Acta Psychologica, 204, 103016.w

Zheng, J., & Meister, M. (2025). The unbearable slowness of being: Why do we live at 10 bits/s?. Neuron, 113(2), 192–204.

